# Design Principles for Autonomous Illumination Control in Localization Microscopy

**DOI:** 10.1101/295519

**Authors:** Marcel Štefko, Baptiste Ottino, Kyle M. Douglass, Suliana Manley

**Author notes:** Authors have made equal contributions (Marcel Štefko, Baptiste Ottino).

## Abstract

Super-resolution fluorescence microscopy improves spatial resolution, but this comes at a loss of image throughput and presents unique challenges in identifying optimal acquisition parameters. Microscope automation routines can offset these drawbacks, but thus far have required user inputs that presume *a priori* knowledge about the sample. Here, we develop a flexible illumination control system for localization microscopy comprised of two interacting components that require no sample-specific inputs: a self-tuning controller and a deep learning molecule density estimator that is accurate over an extended range. This system obviates the need to fine-tune parameters and demonstrates the design of modular illumination control for localization microscopy.

---

Single molecule localization microscopy (SMLM) is a suite of techniques for super-resolution fluorescence imaging that has generated great interest for bioimaging applications. Of these techniques, photoactivated localization microscopy (PALM) and stochastic optical reconstruction microscopy (STORM) achieve super-resolution by exploiting optically-induced transitions of single fluorescent markers between emitting and non-emitting states [1–3]. Due to the tradeoff between spatial and temporal resolutions, there is significant interest in improving their throughput; a single image typically takes minutes to acquire. Several approaches to this problem have been taken including high frame rate imaging [4], tailored illumination for large fields of view (FOVs) [5–7], and automation [8–13]. All of these approaches may be interpreted as means to collect more data at a fixed cost to the microscopist’s time. Automation is a particularly appealing line of technology development because it transfers repetitive tasks to a computer. Automated illumination strategies have enabled multiple field of view acquisitions without user interference, although thus far they have required parameter tuning and *a priori* knowledge of the sample response [8], and have often been targeted to specific samples [9–11].

Automation also has the potential to enable—in real-time, and for a given sample—the optimization of acquisition parameters to produce datasets of the best possible resolutions; that contain a minimal degree of artifacts; and that take the least amount of time to acquire. For example, a suboptimal transition rate between fluorescence emitting and non-emitting states will result in measurements that are either needlessly long or produce artifacts [14–16]. These artifacts must then be corrected in post-processing, lest they lead researchers to incorrect conclusions. Minimizing these errors at the point of acquisition is therefore an important step in the quality control process. Even a well-trained microscopist is unlikely to consistently find the best illumination conditions for every experiment.

In this work, we go beyond specific implementations to address the problem of designing autonomous illumination control systems for adaptive corrections to the active emitter density in any STORM/PALM experiment. We begin by establishing the primary components within the negative feedback loop that comprises the control system. The function of each component is decoupled from the others, which allows us to address their design independently of the system as a whole. With this philosophy in mind, we then develop a new algorithm for each component, intended to reduce the number of user inputs and increase the generality of the overall control system. The first is a parameter-free algorithm for counting emitters in an image. This algorithm, called Density Estimation by Fully Convolutional Networks (DEFCoN), outperforms fluorescent spot counters that are based on matched filters by greatly reducing their bias when signals from individual emitters are highly spatially overlapping. Furthermore, it can be readily adapted to new classes of datasets by re-training the network. The second component is a self-tuning controller that automatically adapts its gain parameters to each specific field-of-view by measuring the fluorescence excitation step response prior to acquisition.

Finally, we reintegrate these components into the control system and show how they may achieve minimal artifacts for PALM/STORM, requiring as input only a single, sample-independent parameter that is general to the problem. These tools are freely provided to the community as the Automated Laser Illumination Control Algorithms (ALICA, pronounced *ah-LEETZ-uh*) plugin for Micro-Manager [17], a free and open-source software library for microscopy acquisition control.

## 1. Design of the Illumination Control System

### A. Optical Control of the Active Emitter Density

Common photodynamical models employed in PALM/STORM are based on a system of states that correspond to the distinct energy levels of a fluorescent molecule. A transition from one state to another during a given time interval is a random event and occurs with a probability that depends only on the rate coefficient ascribed to that transition. In general, the rate coefficients can depend on a number of sample-specific factors, such as a fluorophore’s local environment and its chemical structure. At least one rate coefficient between fluorescence emitting and non-emitting states is proportional to the irradiance (power-per-area) integrated across the fluorophore’s absorption cross section. In direct STORM, for example, the irradiance of excitation light determines the transition rate from the emitting singlet state to the non-emitting triplet state; the irradiance of ultraviolet (UV) light influences the return rate from a non-emitting reduced state to the singlet state [18]. As another example, many photoswitchable fluorescent proteins (PS-FPs) are irreversibly switched into a red-shifted emission state at a rate that depends on the local UV irradiance [2,3].

The existence of these light-induced transitions allows the microscopist to optically tune the density of fluorophores in the emitting state by adjusting the power of the light source(s). Typically, the goal is to adjust the power until there is approximately one active emitter per diffraction-limited area on average. At lower densities, the acquisition will take longer to sample the underlying structure; at higher densities, artifacts begin to appear in the final dataset because localizations may correspond to the centroid of multiple overlapping emitters and not their individual locations.

### B. Design of the Control System

The purpose of the illumination control system is to find and maintain a fixed density of active emitters throughout an acquisition. The system is implemented as a negative feedback loop (Fig. 1).

**Figure 1.**
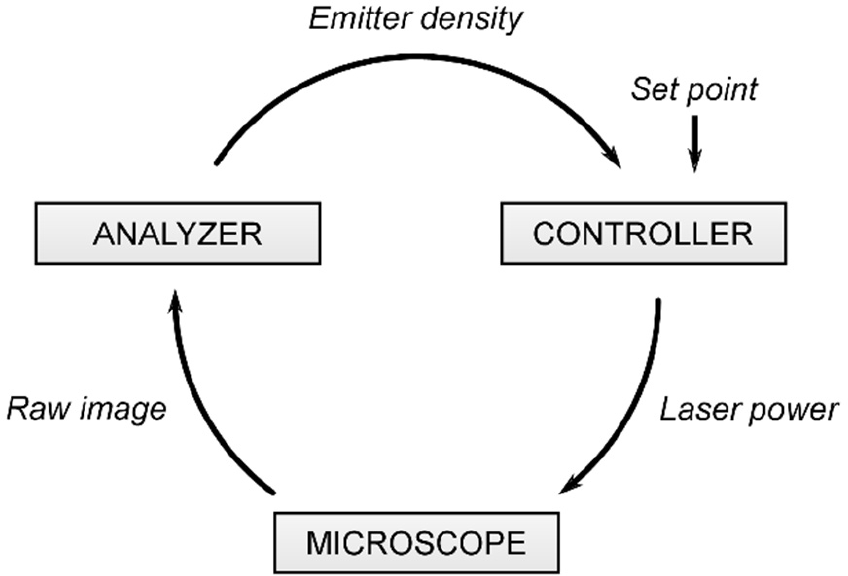
The autonomous illumination control system. The three primary components in the feedback loop are represented as modular blocks, and the data passed between the components are indicated in italics.

The system consists of three generalized components. The first is the microscope, which contains the illumination source and provides raw images from its camera. The images are fed sequentially into an analyzer whose job is to estimate the density of active emitters. (In general, the analyzer may produce estimates of other quantities as well, such as the integrated intensity.) The controller is the third component and takes as inputs the analyzer’s most recent estimate of the active emitter density and the density set point, i.e. the desired emitter density to maintain during the experiment. The controller’s purpose is to compute the power of the illumination source that minimizes the absolute difference between the estimate and the set point. If the difference deviates from zero—as it would at the very start of a measurement or over time due to photobleaching—then the controller will apply a corrective adjustment to the light source’s output power.

The division of labor between the components carries several advantages. The components are weakly coupled; thus, if any component fails to perform its computation before the previous one in the feedback loop, the other components can still continue their work unimpeded. Furthermore, the algorithms for the different components’ functionalities can be exchanged at will without affecting the others. This allows microscopists to adapt the control system to their particular samples and use cases. This also means that the optimization of each component may be seen as its own independent problem, rather than one of the control system as a whole.

In what follows we will leverage these abstractions to independently address a problem of analyzer design and a problem of controller design.

## 2. Estimating Emitter Counts with Density Maps

### A. The Spot Counting Problem

A natural choice for an analyzer for localization microscopy is a module that estimates emitter density. This can be translated into the spot counting problem, which refers to the process of algorithmically counting the number of single fluorescent molecules in an image (Fig. 2). This figure shows a single frame from a simulated PALM acquisition with the ground truth positions of emitters marked as red x’s. The most straightforward way to count the number of spots in this image is to run a spot detection algorithm and simply count the number of detections. Such an algorithm usually involves two steps. First, the image’s power spectrum is whitened and convolved with a filter that amplifies the signal from the fluorescent molecules while simultaneously suppressing the background. The convolutional kernel is often a matched filter whose frequency response is the conjugate of the Fourier transform of the microscope point spread function (PSF) [19]. Small regions of interest (ROIs) surrounding local maxima in the filtered image are then identified as single emitters. The detections in Fig. 2—marked as cyan circles—were identified using a wavelet-based matched filter coupled with watershed segmentation and followed by a calculation of the centroid of connected components as described in [20] and implemented in the software package ThunderSTORM [21].

**Figure 2.**
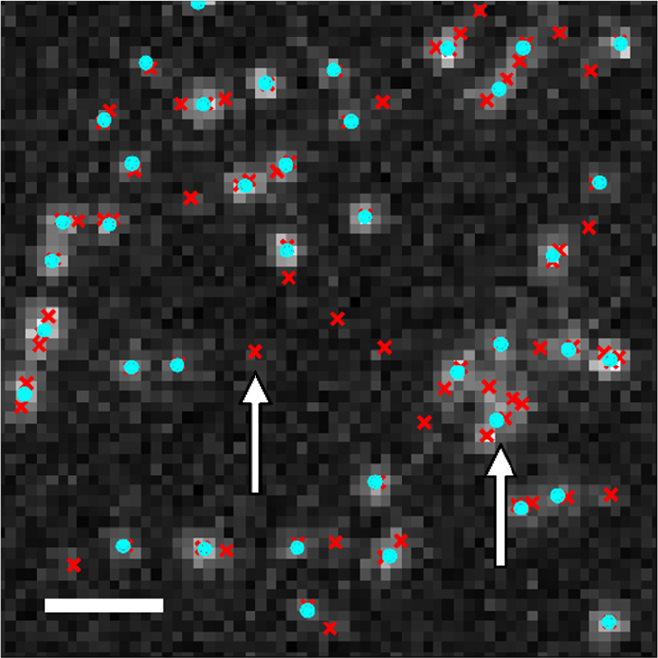
The spot counting problem. A single image from a simulated PALM acquisition demonstrates two types of counting errors: undercounting due to poor signal-to-noise (left arrow) and undercounting due to overlapping PSFs (right arrow). Red x’s: ground truth emitters; cyan circles: detected molecules using a wavelet matched filter. Scale bar: 1 µm.

One can immediately see that the number of detections is less than the number of actual emitters. Furthermore, the bias towards undercounting appears to be a general feature of counting by direct detection and becomes worse as the true density of emitters increases; this makes the spot counter’s response to the true density nonlinear, which is an important consideration in control systems design (Fig. S1). Two types of error contribute to this bias. The first is missed detections due to a poor signal-to-noise ratio (SNR); this error is largely inconsequential because the absence of emitters with poor SNR from a dataset typically does not have an adverse effect on the SMLM reconstruction. The second error arises from the overlapping signals from closely-spaced emitters. In the context of automation performance, this would result in the control system erroneously concluding that there are fewer active emitters than there are in reality, thereby preventing the system from taking corrective action. The nonlinear response of the detection-based spot counter furthermore complicates the controller design because using it may necessitate “gain scheduling,” a process of retuning the control parameters for different densities.

One alternative to spot counting is to compute a quantity from the images that is somehow proportional to the number of spots, such as the sum over pixel values or the time that a pixel value spends above a given threshold [22]. While this approach can be made linear in the number of active emitters, it is susceptible to other types of errors that limit its use to samples where the only significant source of light is from the target fluorophores. Autofluorescence, contaminants, and out-of-focus fluorescence would all bias the emitter count estimate.

Another alternative would be to use multi-emitter subpixel localization routines. (An extensive and recent list of such routines may be found at [23].) In principle, these algorithms can perform unbiased spot counting in the case of overlapping signals by fitting the photon count distributions to models containing multiple emitters. They often require extensive parameter tuning, however, and are too slow to use for real-time applications or large FOVs.

### B. Density Map Regression

The consistent bias of detection-based counters suggests that a new approach is required to alleviate these issues. We therefore reformulated the problem of fluorescence spot counting as a regression problem over a density map (Fig. 3a). In this formulation, a model is constructed that transforms an input image of fluorescent spots into a density map, i.e. a 2D image of the same size as the input and upon which a normalized Gaussian kernel is placed at each ground truth active fluorophore position. The integrated sum of the density map pixels over a subregion is equal to the number of spots it contains; the integral over the full density map is the estimated number of spots within the FOV. Density map regression has been successfully applied to problems in counting pedestrians, cars, and cells [24–26].

**Figure 3.**
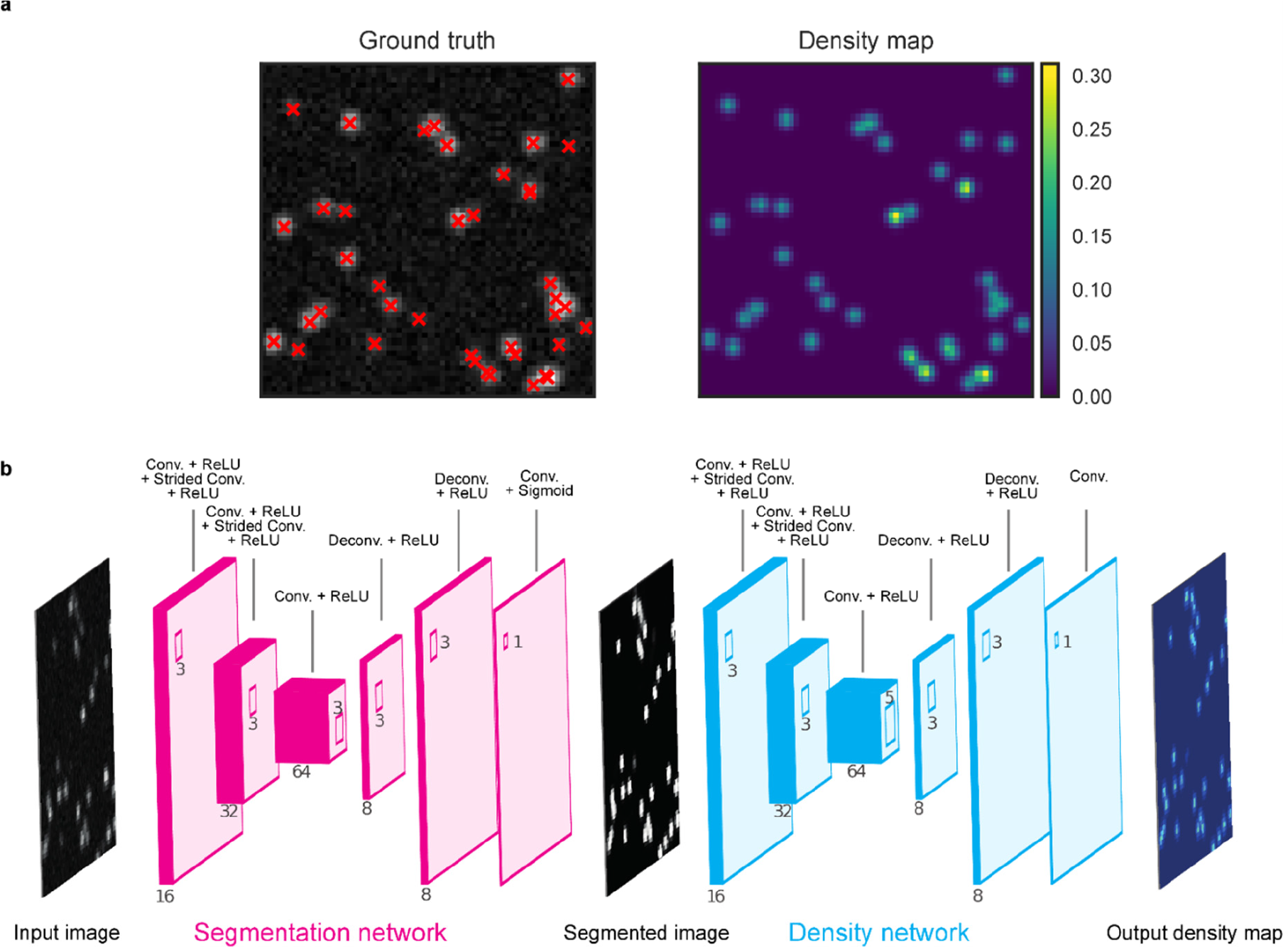
Density map estimation for fluorescence spot counting. a) A target density map generated from ground truth simulated data. The integral over the density map is the number of fluorescent spots in the FOV. Red x’s denote ground truth positions. b) The architecture of DEFCoN.

Previous models for density map regression have utilized an ad-hoc MESA distance [24] refined with ridge-regression [25], a random forest with hand-crafted features [27], or fully convolutional neural networks (FCNNs) [26,28,29]. FCNNs are particularly attractive because they do not require hand-crafted features and the model can be trained directly from images. In addition, their computational complexity scales linearly with the number of pixels, rendering them useful for real-time computation and competitive in terms of speed with detection-based algorithms [30]. To this end, we designed a spot counter called Density Estimation by Fully Convolutional Networks (DEFCoN) for density map regression of images of fluorescent spots (Fig. 3b).

### 1. Network Architecture

DEFCoN’s architecture consists of two fully convolutional networks in series: a segmentation network and a density network. Each network is comprised of layers of (de)convolutional operations and nonlinear image transforms that first form a downsampling path and then an upsampling path. In the downsampling path, the convolutional operations serve to extract features in the image at length scales that increase with each successive layer. In the upsampling path, an output image (either a segmentation map or density map) is constructed from these features. Training DEFCoN means finding the values of all the square (de)convolutional filters that produce accurate segmentation and density maps.

The segmentation network computes a parameter-free segmentation of the input image (Fig. S2). The downsampling path of the segmentation network consists of three convolutional layers with 3 × 3 pixel kernels and separated by strided convolutional layers. The receptive field of the deepest layer corresponds to a 12 × 12 pixel region on the original image, which is large enough to capture information about the shape of clusters of fluorescent spots yet small enough to maintain the speed of the network’s computation. The upsampling path is made of two layers of eight deconvolution kernels followed by a 1 × 1 convolutional layer with a sigmoid activation function. During training, the network’s output is compared to a binary, ground truth segmentation mask using pixel-wise binary cross-entropy as the loss function (Fig. S2). Essentially, the segmentation network performs a per-pixel classification where the output is a map indicating the probability that the value of a pixel is determined at least in part by a fluorophore.

The density network transforms the segmentation map into the final density map and possesses a similar architecture as the segmentation network (Fig. 3b). The deepest convolutional layer is made of 5 × 5 pixel kernels—making the receptive field 15 × 15 pixels—and the final layer has a linear, rather than sigmoid, activation function.

The reason for the inclusion of the segmentation network in DEFCoN is empirical; we found that the density estimation network alone does not generalize well to new datasets. This is likely due to the large degree of similarity between the input images and the density map estimates. In the absence of the segmentation network, rather than learning meaningful representations of what a fluorophore looks like, the density network would instead learn how to minimize the counting error through subtle pixel-wise transformations. The result is that non-zero values would be sporadically placed in the background pixels of the resulting density maps, significantly biasing local counts. The addition of the segmentation network is our solution to avoid fine-tuning for improved generalization, such as is done in [26].

### 2. Training DECON

The DEFCoN network is trained in two phases (Fig. S2; Supplement 1). The segmentation network is trained alone in the first phase using ground truth segmentation masks generated from simulated data. Next, its weights are frozen and the combined segmentation/density network is trained in full, this time with ground truth density maps also generated from simulated data. As in [28], the loss function that is used for backpropagation while training the full network is comprised of two terms.

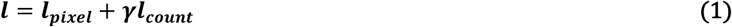

The first term is simply the sum of the squared pixel errors

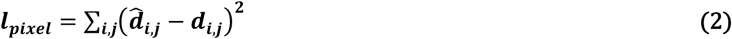

where 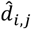 and *d*_*i,j*_ are the values of pixel *i, j* in the predicted and ground truth density maps, respectively. The second term, *l*_*count*_, penalizes the network for counting the number of spots incorrectly. Since the count is merely the sum of all the density map pixel values, this term is expressed as

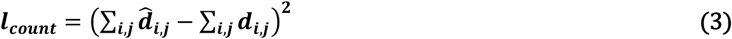

The parameter γ varies the relative weight attributed to each term. If *γ* is too small, each pixel can adopt a small offset that leads to a systematic counting error in the density map; if *γ* is too large, the network will lose some local information, resulting in misshapen kernels. We empirically found a value of *γ* = 0.01 to give the best results. For a more detailed description of how the network was trained, please see Supplement 1.

### C. DEFCoN Performance

We tested DEFCoN against the ThunderSTORM implementation of the wavelet filtering and watershed algorithm because it is currently one of the top-performing segmentation algorithms for SMLM and performs well when spots weakly overlap [20,21]. We first generated several simulated SMLM stacks consisting of 100 images, 128 × 128 pixels in size, of randomly distributed fluorophores with different mean densities of active emitters (in units of *µ*m^-2^) and SNRs (Fig. S3). Here, the SNR is defined as the ratio between the maximum value of the pixels spanned by the image of a single fluorescent molecule and the standard deviation of the neighboring background pixel values. Next, we applied each algorithm to the SMLM stacks and calculated a performance metric to compare the two, in this case the counting error:

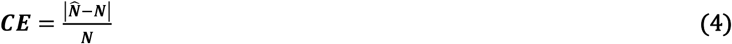

where 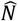 and *N* are the predicted and ground truth fluorophore counts, respectively. The mean counting errors from the test are displayed in Fig. 4.

**Figure 4.**
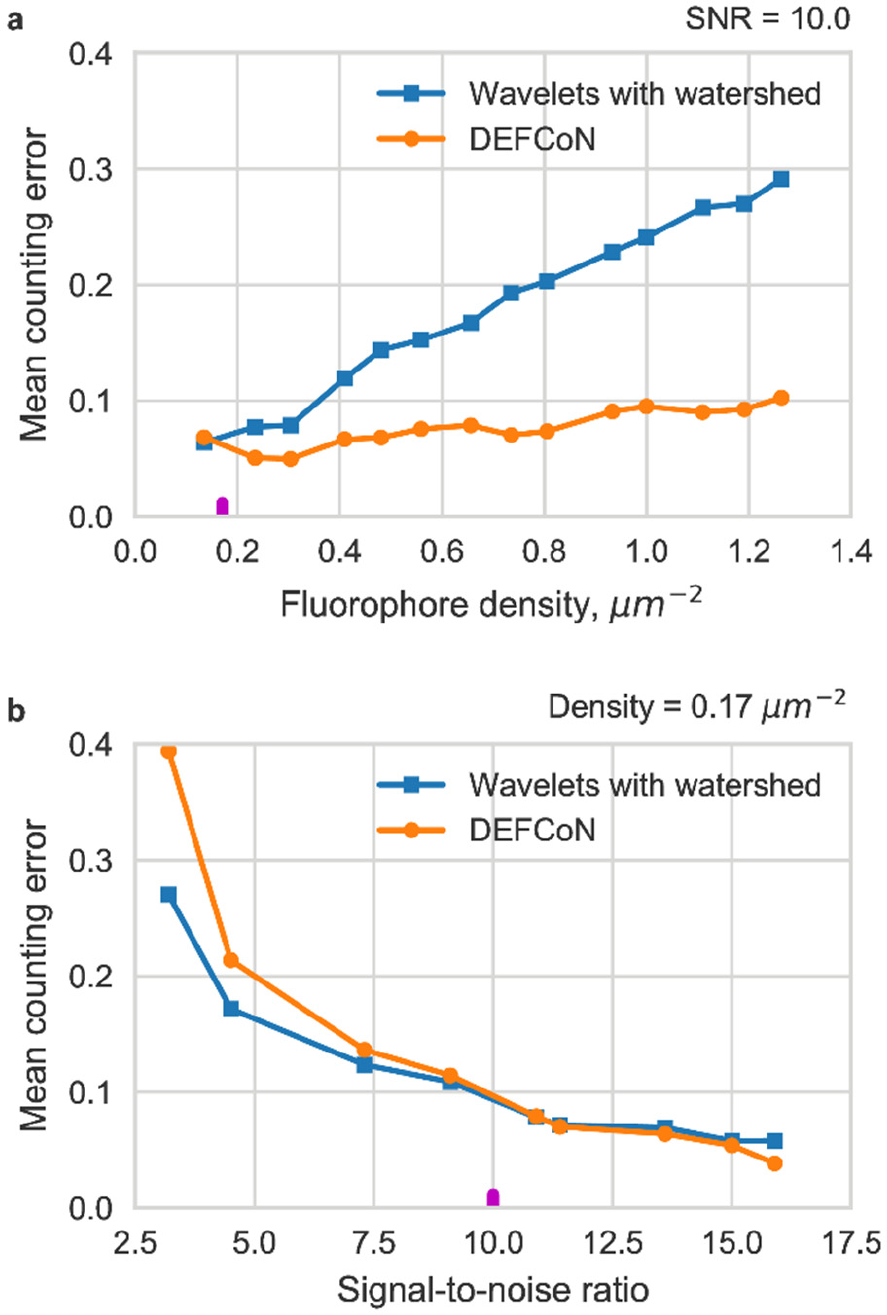
Comparison between DEFCoN and the wavelet filtering/watershed method from [20]. a) The mean counting error’s dependence on the density of randomly distributed fluorophores. The magenta tick indicates the density in the simulated datasets for panel b. b) The dependence of the error on the SNR. The magenta tick indicates the SNR of the simulated datasets in panel a.

DEFCoN performs extremely well at counting fluorescent spots across a range of fluorophore densities (Fig. 4a), with the mean counting error increasing from approximately 0.07 to 0.10 for sparse to moderate densities of randomly distributed emitters. The wavelet filtering algorithm with watershed performs as well as DEFCoN at sparse densities. However, its mean counting error grows linearly with density at a rate that is ~5-7 times faster than DEFCoN’s for an SNR of 10. DEFCoN performs slightly worse than the wavelet/watershed method at low SNRs (Fig. 4b), with a mean counting error of 0.39 for DEFCoN vs. 0.27 for wavelets. This disparity decreases with increasing SNR until they perform similarly above an SNR ~ 7. In addition, we qualitatively compared DEFCoN, wavelet filtering/watershed in ThunderSTORM, and a simple spot counter based on local maxima identification [31] by running each on a simulated microtubule dataset (Fig. S4). The results show the decreased bias of DEFCoN relative to the other two methods. Though the bias has not been entirely eliminated, the linear response regime of DEFCoN covers a significant proportion of the full range of densities applicable to SMLM.

Equally important for real-time spot counting is the speed with which each algorithm executes. A rough criterion is that the time required for the algorithm to produce a spot count should be less than 10 ms, which is the fastest running exposure time of commercially available sCMOS cameras with a 2048 × 2048 pixel ROI. (EMCCD cameras are slower than their sCMOS counterparts at the full ROI size and equivalent bit depth.) The results of the speed comparisons calculated from the same dataset are shown in Fig. 5.

**Figure 5.**
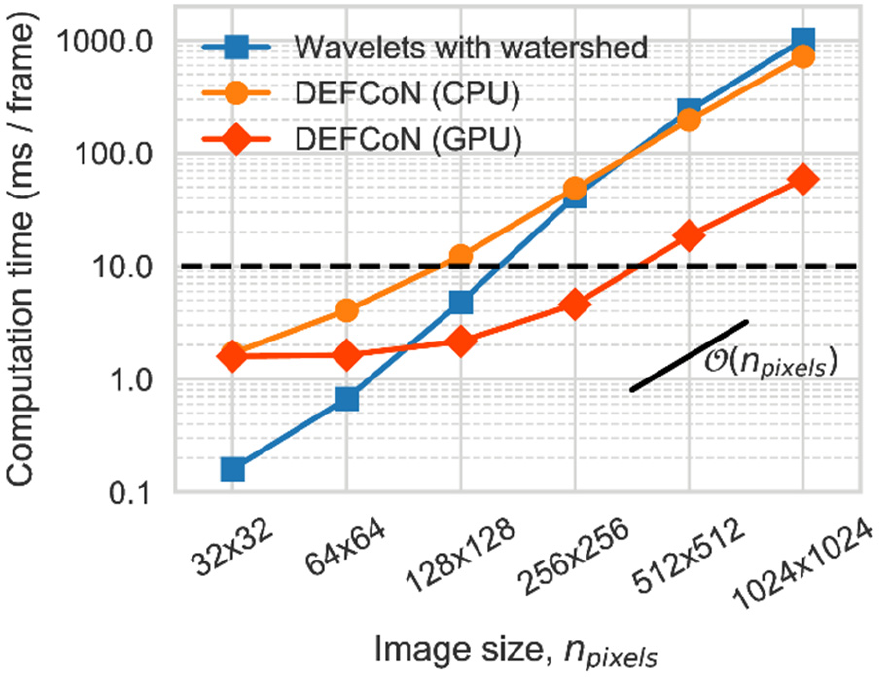
Execution times for DEFCoN and wavelet-based segmentation. The dependence of the execution time on the image size scales linearly with the number of pixels.

When implemented on a GPU, DEFCoN can produce a spot count in less than 10 ms for images that are 512 × 512 pixels in size. The CPU implementation of DEFCoN performs similarly in speed to a CPU-based wavelet/watershed combination for image sizes larger than 256 × 256 pixels. As expected, the computation time of DEFCoN grows linearly with the number of pixels and is independent of the density of fluorescence spots (Fig. 5).

Finally, we tested DEFCoN on the RealLS and RealHD datasets from the 2016 SMLMS Challenge [32]. RealLS is a low density dataset where the active emitters are sparsely distributed in space; RealHD is a high density dataset containing many overlapping fluorophores. Because no ground-truth data exists for these experimentally-derived datasets, 10 frames from RealLS and 5 from RealHD were given dot annotations by hand, where each dot marked the ground truth position of a visible fluorophore (Fig. S5). The mean counting errors for DEFCoN and wavelets/watershed are displayed in Table 1. As expected, both algorithms perform well at low density, but DEFCoN produces more accurate counts (with respect to the annotations) in the high density dataset.

**Table 1.** Mean counting errors on real datasets.

|  | RealLS | RealHD |
| --- | --- | --- |
| DEFCoN | 1.4 | 2.4 |
| Wavelets/Watershed | 1.7 | 19.0 |

Taken together, these results indicate that DEFCoN outperforms the state-of-the art detection-based approaches for fluorescence spot counting. The improved linearity and ability to work across a large range of active emitter densities makes its application in an illumination control system both general and robust.

## 3. Controller Self-Tuning

### A. The Controller Tuning Problem

Having dealt with the problem of making accurate estimates of the density of emitters, we now turn to the problem of control: how does one compute the required illumination intensity to activate and maintain a set density of emitters? In what follows, we will restrict the discussion to control of a UV illumination source because fluorophores respond strongly to even relatively weak UV irradiance and because it controls the density of active emitters in both PALM and STORM.

We can divide a PALM acquisition into two distinct periods for which we would like the controller to autonomously determine the optimal laser power. During the initial period at the very beginning of the acquisition, the problem is to determine how much power activates the ideal number of emitters. During the second period, which extends from when the emitter density has reached a quasi-steady state to the end of the acquisition, the problem is to make continuous and small adjustments to the power to compensate for a gradual decline in the active emitter density due to photobleaching.

Automation methods for photobleaching correction were first proposed in [8] and [22]. The controller in [8] counted fluorescent spots in real-time using the wavelet approach of [20], accepting as inputs three free parameters. The first two are thresholds for the average and maximum emitter counts; if either of these two values exceeds ±15% of their original value, then the controller adjusts the illumination power. The third parameter is the amount by which the illumination power is adjusted during each step. Likewise, the controller of [22] also makes discrete adjustments to the illumination power in values that are predetermined by the user. In addition, it requires a threshold value to separate the noise from the signal and another threshold that helps identify and remove pixels from the analysis that are always active, such as those contaminated by autofluorescence from dust particles.

Parameter tuning for these control systems may be performed in exploratory experiments to collect *a priori* knowledge about the typical sample response. The system performance will necessarily depend on how well the parameter values generalize to variability in the response during and between acquisitions. Large variability in labeling density, sample preparation, and the appearance of edge cases like brightly autofluorescent dust particles can invalidate a previously defined set of control parameter values, resulting in a suboptimal density of activated fluorophores.

To our knowledge, no one has yet addressed the first problem, which is determining the optimal control parameters to use on a previously unseen FOV. We therefore addressed these issues by implementing a self-tuning controller that adapts itself to each FOV.

### B. Self-Tuning for Density Set Point Control

In the context of SMLM, our strategy is to implement a proportional-integral (PI) controller (Fig. 6) to compute the power of the UV illumination source that will maintain a constant density of active emitters. It accepts two inputs: the estimate of the density of emitters *N(t)* and the desired density *N*_*0*_, which is also called the set point. The difference between these two quantities is the error signal *e(t)* = *N(t) - N*_*0*_ which is fed in parallel into the proportional and integral block components. The computed power *P* of the illumination source is

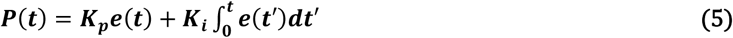

where *K*_*p*_ and *K*_*i*_ are the proportional and integral gain, respectively. The value in choosing PI control over either purely proportional control or stepping the illumination by pre-determined amounts is that it can maintain a long-term zero error signal while still achieving a fast response to perturbations [33].

**Figure 6.**
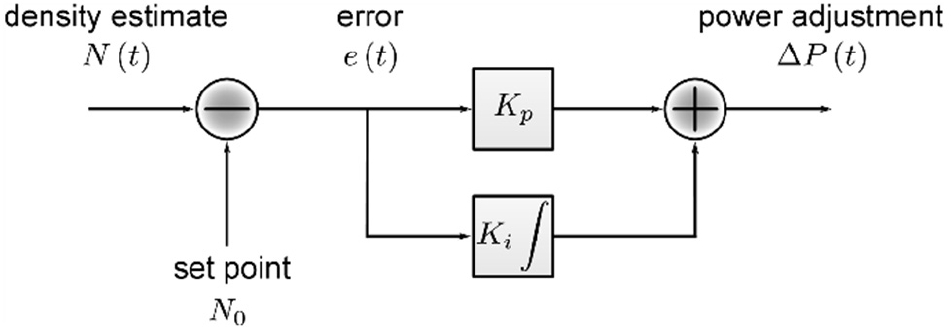
A proportional-integral (PI) controller.

Finding the correct values for the gain parameters is essential to achieving a stable and fast response to changes in both the set point and error signal. Severe oscillations in the illumination output or a slow response to changes in the emitter density may occur when the PI controller is not properly tuned. Furthermore, the optimal values for the gain parameters will vary with each FOV. For these reasons, we implemented a self-tuning procedure that is based on a set of rules derived from internal model control known as lambda tuning [34]. These rules are used to calculate the optimal values for *K*_*p*_ and *K*_*i*_ on a given FOV by measuring the sample’s step response to UV light (Fig. 7).

**Figure 7.**
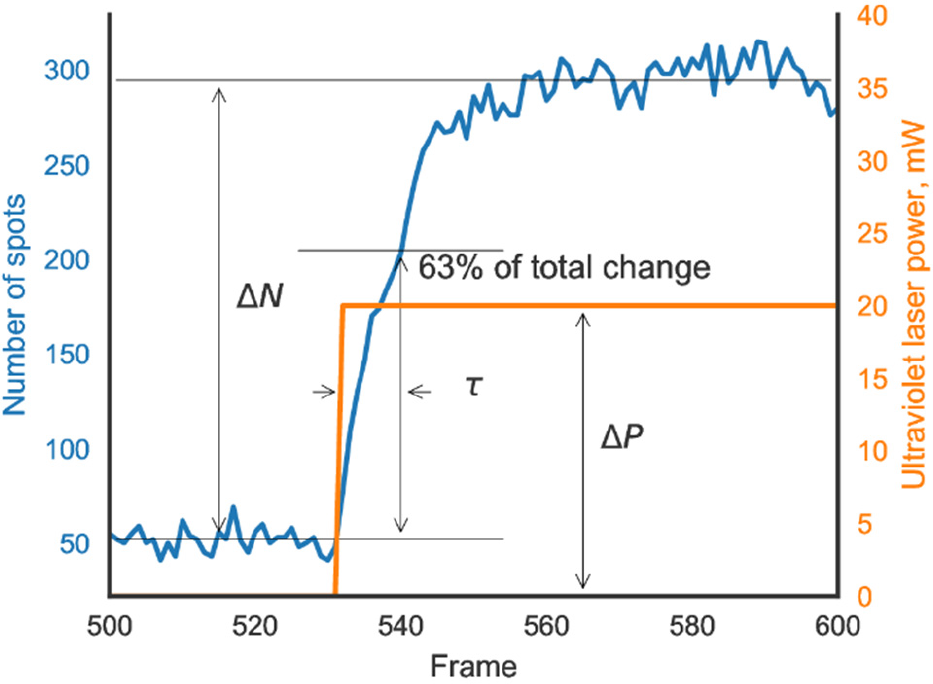
Construction of the self-tuning procedure for the PI controller.

The lambda tuning rules for the PI controller are

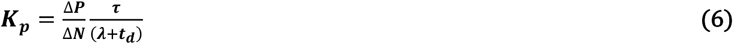

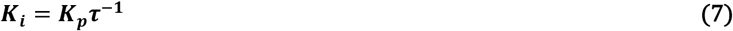

In these expressions, Δ*N* represents the change in the analyzer’s average output in response to a step change Δ*P* in the output power of the illumination source. τ and *t*_*d*_ are the response time and the dead time of the system, respectively. The former represents the amount of time it takes for the analyzer’s output to reach approximately 1 - *e*^-1^ ≈ 63% of its quasi-steady state value at the new laser power, whereas the latter represents the time after the change in laser power when a response is first detected. In our experience, the initial change in emitter density is nearly instantaneous when compared to the exposure time for a camera frame, so we set *t*_*d*_ to zero. Small values of the parameter *λ* will result in a fast response to changes about the set point, whereas large values will result in a slow response. The lambda tuning rules produce a response to a change in set point that settles out in a time of approximately 4*λ* without overshooting the set point value. *λ* = 3τ is recommended for stable set point control [34].

In practice, we find that it is easy to measure the step response *∆N* but difficult to precisely measure τ. Fortunately, the value selected for τ varies little between experiments and need not be precise; a guess often suffices. For example, τ ≈ 10 frames and *t*_*d*_ = 0 frames in Fig. 7. Using the recommended value from the lambda tuning rules of *λ* = 3τ = 30 frames, this means that *K*_*p*_ ≈ 0.33 *∆P⁄∆N*—irrespective of the value of τ—and *K*_*i*_ = (0.03 frames^-1^) ∆*P*/∆*N*. Even if τ was overestimated by an order of magnitude, the controller would bring the system into a quasi-steady state within a small fraction of the total acquisition time, which often extends over tens of thousands of frames.

When applied at the beginning of an acquisition, the self-tuning procedure determines the gain parameter values that map the emitter density to the laser power. This circumvents the need to “search” for the correct power in step-wise increments. There remains, however, another problem that results when the integral term in Eq. 5 becomes large. The integral acts as a form of memory for the error signal and, if allowed to accumulate, can cause the controller to become saturated such that it only outputs its maximum or minimum value. Large error signals accumulate in the controller’s memory, for example, when selecting a value for the set point that is either too high for the available illumination power or close to zero. (In the latter case, spurious detections keep the value of the measured density above zero and cause the integral term to accumulate a negative value.) This condition, more generally known as integral windup [33], is solved by placing upper and lower limits on the value for the integral term. The upper limit is set as the difference between the maximum possible laser output and the value of the proportional error term; the lower limit is the difference between the minimum output and the proportional term.

The last remaining piece of the controller is the determination of the set point, which is addressed in the next section.

## 4. Set Point Determination

### A. The Problem of Optimal Set Point Determination

In the previous discussion, we took for granted that the optimum value for the set point could be easily determined. Roughly speaking, the optimum density of emitters should produce super-resolved reconstructions with the fewest artifacts, the best resolution, and should take the least amount of time. However, the autonomous determination of an optimum value that satisfies these requirements is more challenging than it might first appear. For one, the optimum density is not a global property but varies across the FOV with the structure’s dimensionality [15]. Second, the quality of a super-resolution reconstruction depends on the algorithm chosen to perform the reconstruction [35]. The control system would therefore have to decide whether single or multi-emitter fitting routines would be more suitable and activate the required density accordingly. Third, we lack a rigorous, functional definition of SMLM image quality for real-time optimization. In part, this is because a SMLM image is a function of all the images that contribute to the eventual reconstruction; it is difficult to predict the final image quality as the data is being acquired. Finally, the tradeoffs that one is forced to make between the number of artifacts, resolution, and acquisition time make this a multiobjective optimization problem whose general solutions contain not one but families of so-called “Pareto optimal” solutions. To select one solution from this set requires that the microscopist explicitly specify the degree of tradeoffs that she or he is willing to make between these quantities.

For these reasons, we have instead employed a heuristic solution to the problem: the criterion that there should not be more than one emitter active per diffraction-limited area in any given camera frame. (We note, however, that other heuristics may be incorporated into this modular framework.)

### B. Maximum Local Count Control

The approach we employ here is to compute the highest local density of active emitters in an image from a density map estimate and subsequently use this quantity as the controller’s input. At the same time, we choose as the set point a free parameter that is slightly higher than one emitter per area spanned by the PSF.

The maximum local density of emitters arises naturally from a density map estimate because the sum of the pixels over any subregion produces the number of emitters within that same subregion. We can transform the DEFCoN output into a map of local emitter densities through an extension of the so-called “gliding box algorithm” [36]. Briefly, a kernel of size *n×n* pixels and whose values are all unity is convolved with the density map. During a single step of the convolution, the value of the pixel currently at the center of the gliding window is replaced with the sum of the pixels that fall within the window. The final result is a map whose values represent the local densities of emitters (Fig. S6). The maximum local count (MLC) is the maximum value over the entirety of this new map:

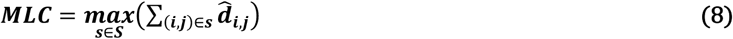

where *S* is the set of all *s* subregions within an estimated density map 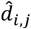.

Due to the stochastic switching of emitters, large fluctuations are expected in the MLC time-trace. A more reliable control value is therefore the average MLC (AMLC), which is simply a moving average over the most recently computed MLC values. (A Kalman filter may be a more robust solution, but photobleaching causes the mean density of emitters to decrease so slowly that a simple moving average will typically suffice.)

Fig. 8 displays the results of activating fluorophores on a simulated 2D microtubule network with different AMLC values. Briefly, fluorophores whose photodynamics followed that of a simple two state ON/OFF model were simulated with a 2D Gaussian PSF. 5000 raw frames were generated for each of four different mean fluorophore off-times and the fluorophores were localized with subpixel accuracies in ThunderSTORM [21]. The AMLC for each stack was computed over 7 × 7 pixel windows (pixel size: 100 nm) from DEFCoN’s density maps.

**Figure 8.**
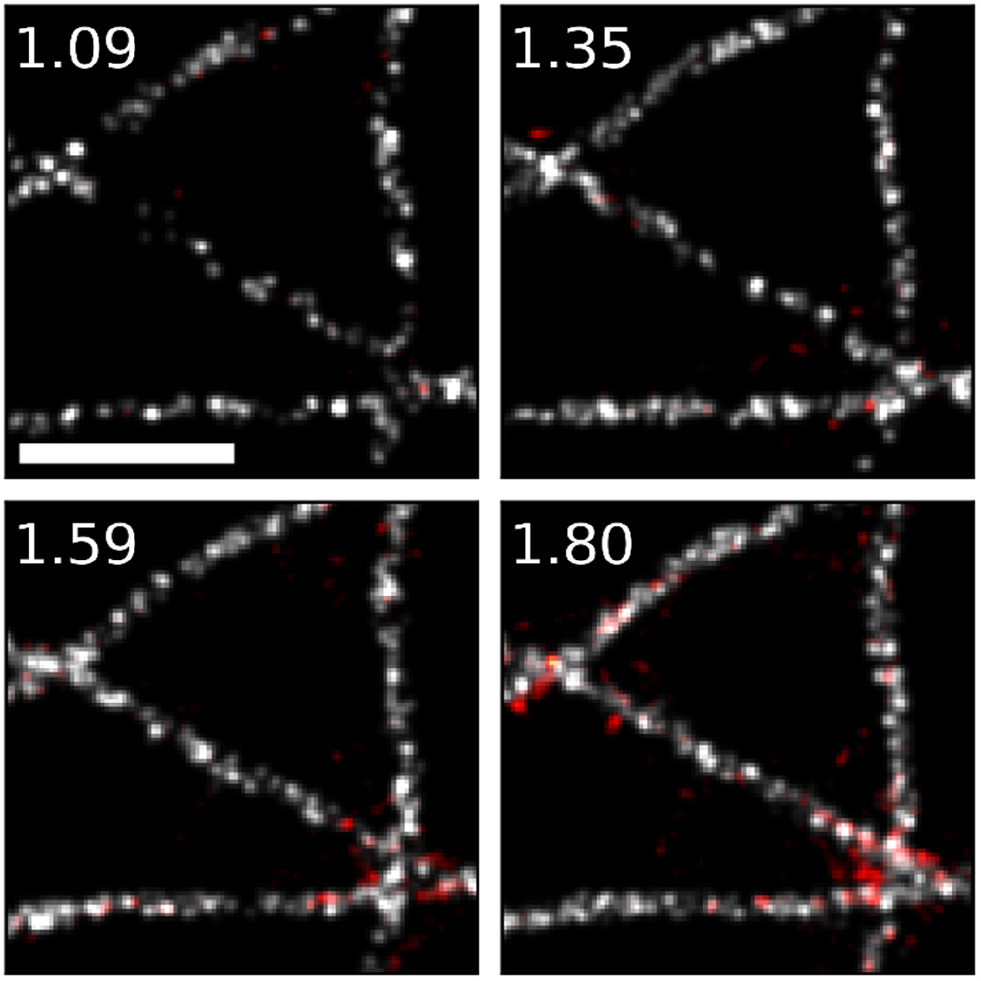
The AMLC as a heuristic for set point determination. AMLC values are in the upper left corner of each image. False positive localizations are in red. Scale bar: 1 µm.

Qualitatively, we see that the AMLC correlates well with the final density of localizations on the structure and therefore the final resolution of the image [37,38]. It is also positively correlated with the number of false positive localizations (red) which reflects, for a fixed acquisition time, the tradeoff between resolution and the degree of artifacts. Importantly, the AMLC is a local property. Regions with the highest ground truth density of fluorophores, such as the intersections of different tubules, will contribute more often to the AMLC than regions with a low ground truth density. As demonstrated in Fig. 8, these high density regions are precisely those that are most prone to artifacts.

In this light, the AMLC is a heuristic for determining a set point that minimizes artifacts across the entire FOV. It does this at a cost of potentially increasing the acquisition time because the regions that are less dense with fluorophores may be under-activated. One possible solution could instead be to average the MLC over time *and* space, taking care to exclude empty regions from the spatial average. The value in this example is that it demonstrates how to apply domain knowledge when determining the set point: translate the set point from a quantity that depends on the FOV into one that is more general to the problem.

## 5. Discussion

To promote further development in this field, we have packaged these tools into a plugin for Micro-Manager 2.0, an open source software suite for microscope control [17]. The plugin, called ALICA, implements the control system in Fig. 1 and allows users to select between different algorithms for the analyzer and controller based on their needs.

One may ask though why control systems like those developed here are necessary for SMLM or more generally for super-resolution. Beyond saving time and increasing throughput, they stand to improve the quality and reproducibility of microscopy data. Properly designed and tested algorithms do not make mistakes. Once established, they do not require laboratory training to distinguish good imaging from bad. For this reason, they can play a significant role in helping to ensure that super-resolved data acquisition is unbiased and contains minimal errors.

It is important therefore to distinguish between automated and autonomous, or “optimal,” imaging. Simple automation of an acquisition implies that microscope hardware executes a predetermined list of actions without regard to the quality of the data. It can save time, but some of its benefits are offset by the rigorous quality control steps that must necessarily follow acquisition. Optimal imaging systems on the other hand attempt to find the best combination of hardware settings for producing datasets that are most faithful to the structure in terms of both accuracy and precision, while simultaneously ensuring maximal resolution and minimal acquisition time. Optimal imaging is much harder than automation to implement. As demonstrated here, it requires engineering multiple components of feedback loops to make them robust against a range of possible sample conditions. It also involves reducing the number of free parameters that require tuning; in doing so we reduce the number of assumptions and instead force the measurement system to adapt to the sample. Finally, with both the self-tuning PI controller and the AMLC we demonstrated the value in constructing systems whose control parameters reflect quantities within the general problem domain, rather than those that are sample-specific.

DEFCoN works by recognizing how to count molecules based on the shapes of multiple, overlapping spots. It does not, however, currently extract count information that is encoded in time. Accounting for such information may improve its accuracy in high density conditions, such as when imaging so-called zero-dimensional structures that appear as diffraction-limited spots in widefield images. We expect that extending DEFCoN to include temporal information would further decrease its bias, although it already works well across the range of emitter densities frequently encountered in SMLM. The real value in incorporating temporal information would be to extend ALICA’s toolset into more general fluctuation-based super-resolution modalities, such as SOFI and SRRF [39,40], which excel in dense environments of active emitters. We also note that the purpose of DEFCoN, i.e. counting, is different from other recent Deep Learning approaches to SMLM, Deep-STORM and DeepLoco [41,42]. DEFCoN directly computes local densities of molecules and is tailored for real-time control systems. Both Deep-STORM and DeepLoco are intended to compute localizations and are tailored for high precision, post-acquisition analysis.

In summary, we presented solutions to three problems in illumination control for localization microscopy. In doing so, we exercised the design principles that we believe are most important for autonomous super-resolution illumination systems: implement feedback, modularize the control loop, optimize the subsystems, and translate control parameters into quantities that are independent of the sample. Finally, we provide open-source tools so that these developments may continue to improve the quality and reproducibility of super-resolution microscopy data.

Data associated with this manuscript is available at http://doi.org/10.5281/zenodo.1212352 [43]. The software tools presented in this work—ALICA, DEFCoN, and SASS—are available at https://github.com/LEB-EPFL.

## Acknowledgment

We thank Frank Scheffold, Ricardo Henriques, Seamus J. Holden, Volkan Cevher, Paul Rolland, Daniel Sage, and Christian Sieben for their generous feedback and fruitful discussions.

See Supplement 1 for supporting content.

## 1. Spot Counter Bias

The bias of spot counters refers to the nonlinear shape and of the curves obtained by plotting the estimated counts against the true number of active emitters in an image. Spot counting by detection with matched filters and local maxima search displays a bias towards undercounting at high densities, indicated by a saturation of the curve.

To demonstrate this effect, we simulated 15,000 frames of a 2D SMLM image stack and ran different spot counting algorithms on the stack for comparison. The average number of active emitters was increased every 1000’th frame by slightly increasing the transition rate from the off to the emitting state. The fluorophores were randomly arranged on a 2D microtubule network from the 2016 SMLMS Challenge [1] and were modeled with a two-state system whose simulated lifetimes were exponentially-distributed. We tested four different algorithms for counting: three detection-based algorithms using filtering and peak finding and AutoLase, an algorithm which indirectly counts spots by measuring pixel on-times [2]. The detection-based counters included SpotCounter [3], a simple local peak finder, the wavelet + watershed method of Izeddin et al. [4], and ComDet, a spot finder optimized for heterogeneous backgrounds [5].

Fig. S1 displays the results of this simulation for varying input parameters of the different algorithms. With the exception of the wavelets/watershed algorithm at the smallest spline scale parameter, all of the response curves display a saturation at high densities within the range tested here. The wavelet/watershed curve that does not saturate, however, grossly over counts the number of spots at low densities because it detects noisy pixels.

The AutoLase tests were performed by simulating 150,000 frames and recording only every 10’th frame to avoid introducing spurious correlations from emitters that were on for multiple frames into the count. The AutoLase curve with a threshold just above the pixel offset value (a threshold of 120 and offset of 100 ADU) displays the best linearity over the range of emitter counts. However, this comes at a cost of not knowing the absolute count numbers but rather the pixel on-times. (Note the vertical offset relative to the gray line.) To avoid bias by bright continuous objects such as dust particles, AutoLase implements an *ad hoc* routine to remove pixels from the analysis that have been on for too long. Furthermore, AutoLase assumes the same photodynamics model that was used in the simulation; it is not clear how well it performs in realistic scenarios where the fluorescence dynamics can no longer be modeled as a memoryless two-state system. Finally, the underperformance of ComDet relative to the other algorithms for counting should not be taken as a negative sign about ComDet’s segmentation performance. Because it combines clusters of spots into single detections, it undercounts by design.

**Supplementary Figure S1.**
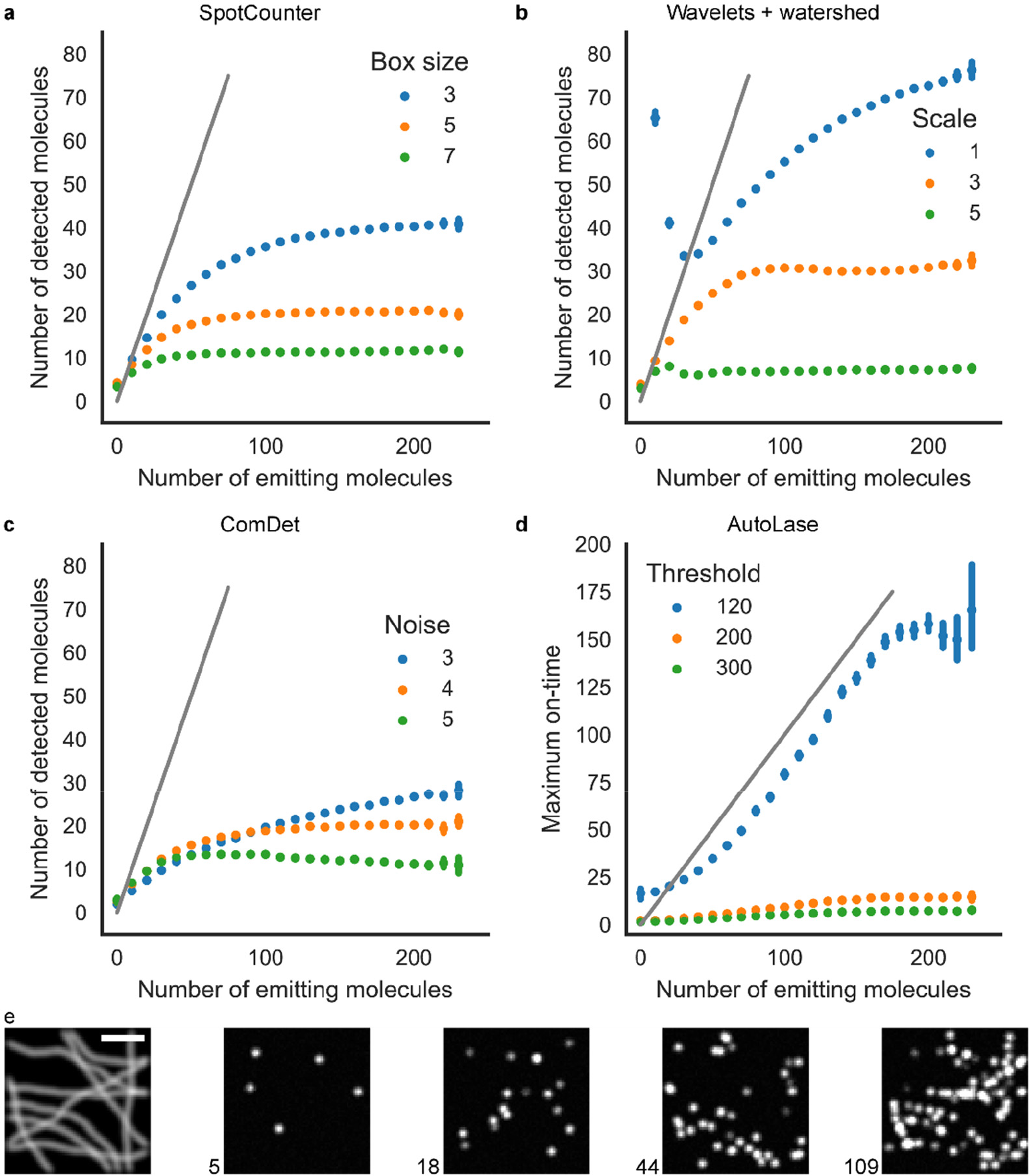
Spot counters and bias towards undercounting. The gray line indicates a perfect, unbiased result. Data points are binned averages with error bars representing the 95% confidence interval of the mean. a) SpotCounter with a variable “Box size” parameter. b) The wavelet/watershed algorithm with a variable B-spline scale argument. c) ComDet with a variable noise setting. d) AutoLase with various threshold values. e) The widefield and example images from the simulated stack with the ground-truth number of active emitters indicated. Scale bar: 2 µm.

## 2. DEFCoN Training

### A. Training data generation, augmentation, and preprocessing

DEFCoN is trained exclusively on simulated data from SASS, our in-house simulation and development platform [6]. The training process is illustrated in Fig. S2. 89,500 64-by-64 pixel training images were generated with signal-to-noise(SNR) varying between 2.0 and 17.2 and with fluorophore densities between 0.0 and 1.5 µm^−2^, to reflect the broad range of conditions encountered in SMLM. The camera pixel size varies between 50 nm and 135 nm. Half the training sets contain out-of-focus fluorophores using the Gibson-Lanni model for the point spread function (PSF) [7,8]. A third of the set is built using realistic microtubule simulations, while the other two are made of randomly distributed fluorophores. Finally, a low-frequency random background is generated on half of the images using simplex noise [9,10].

The generated images are augmented further by random shifts of the brightness and contrast. The network is trained using the full images. Training on random crops of the images was also tested, but this method does not give better training performance. Finally, before being fed to the network, the input images are preprocessed using linear normalization, i.e. histogram stretching. If *I*_*i,j*_ is the original intensity of pixel *(i, j)*, then the transformation is given by:

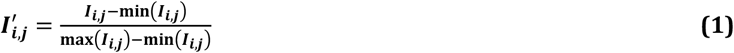

As a result, every pixel value in the image is between 0 and 1. Normalizing the inputs improves training speed and generalization.

**Supplementary Figure S2.**
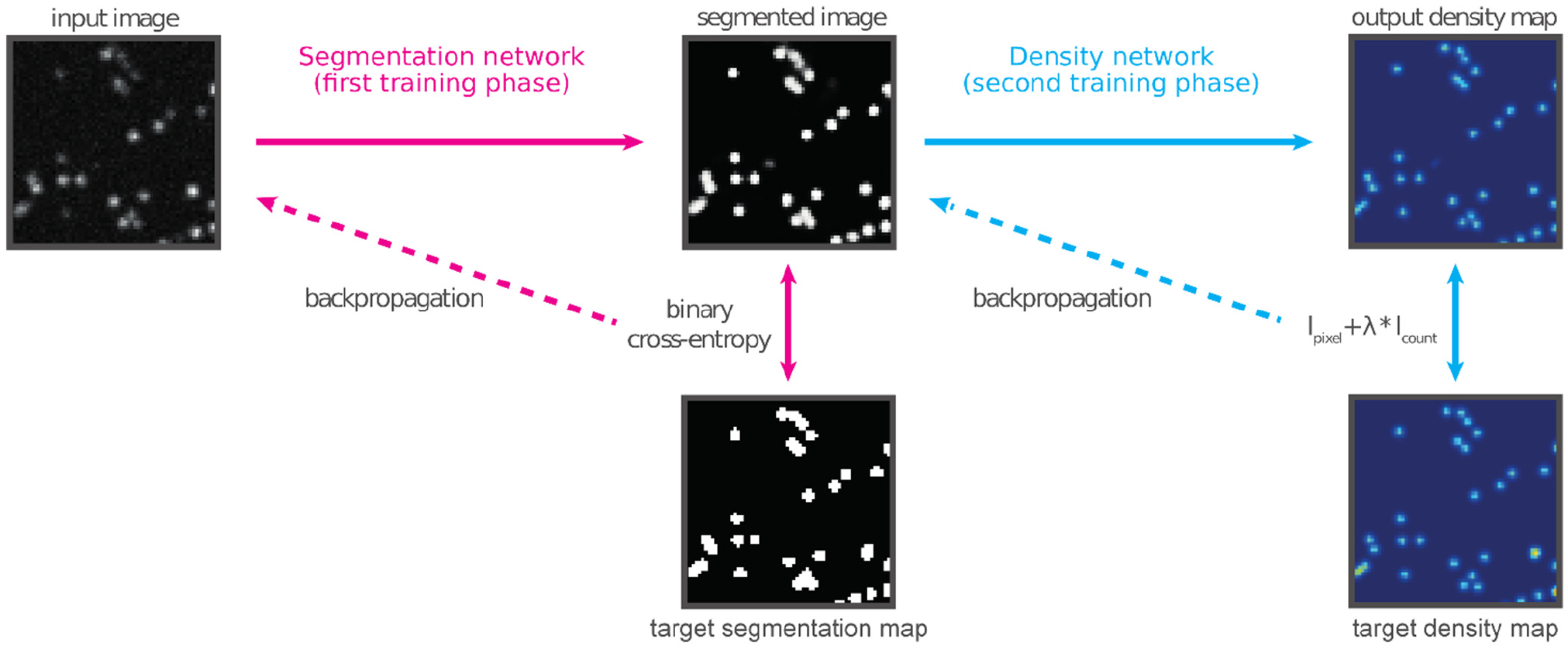
Training DEFCoN’s neural networks. The training takes place in two steps: first the segmentation network alone is trained on target segmentation maps. Then its weights are frozen and the full network is trained on the target density maps.

### B. Target construction and training

To train DEFCoN, two target images were built for each image: one segmentation mask and one density map. The density maps are created by adding Gaussian kernels to an empty image at the ground-truth positions of the emitters. Emitters whose total signals were less than 250 photons per frame were not included in the density map because they were likely to have had a very poor SNR in the simulated image. For the kernels, we use the standard deviation σ = 1 pixel. We found this value to be a good compromise; smaller kernels do not have a good resolution, while larger kernels overlap too much. Each training image is stored with its corresponding density map.

The segmentation masks are built from the density maps. First a threshold is applied to the Gaussian kernels in the ground truth density maps; every pixel with density-map value over 0.03 is given the value 1, every pixel under is set to 0.

Training is done in two phases (Fig. S2). First, the segmentation network alone is trained on 80550 images, holding out 8950 images for validation. A 0.5 dropout regularization layer [11] is applied after the deepest convolutional layer to prevent overfitting. The training is stopped when the performance on the validation data has stopped improving (early stopping). This double regularization (dropout and early stopping) ensures that the network generalizes well to new datasets.

The complete network is then trained end-to-end, feeding the images and comparing the output to the ground truth density maps. However, to keep the segmentation task completely separated from the density map inference, the weights in the segmentation network are frozen; only the weights of the density network are adjusted with backpropagation during the second training phase. In this configuration, the density network is trained with the same validation, optimization and regularization parameters as the segmentation network.

### C. Implementation Details

The network is implemented and trained using Tensorflow 1.3.0 [12] with the Keras 2.0.8 API by François Chollet [13]. We use the Adam optimization algorithm [14], with an initial learning rate of 0.001 for both training phases and with a batch size of 32. These parameters are not of crucial importance, but are chosen to achieve fast convergence. Training takes between one and two hours with an Nvidia GeForce GTX 1060 6GB GPU and an Intel Core i5-7600K CPU, clock speed of 3.80 GHz.

### D. Simulated dataset for comparison

Fig. S3 shows various examples of the dataset of simulated SMLM images that were used to compare the counting accuracy of DEFCoN to that of the approach introduced by Izeddin et al. [4]. These example images were simulated with SASS [6].

Fig. S4 demonstrates the degree of undercounting of three different spot counters: DEFCoN, the ThunderSTORM [15] implementation of the wavelet-based filtering approach of Izeddin et al., and Spot counter [3]. The tests were performed on the simulated dataset whose examples are displayed in Fig. S1e.

**Supplementary Figure S3.**
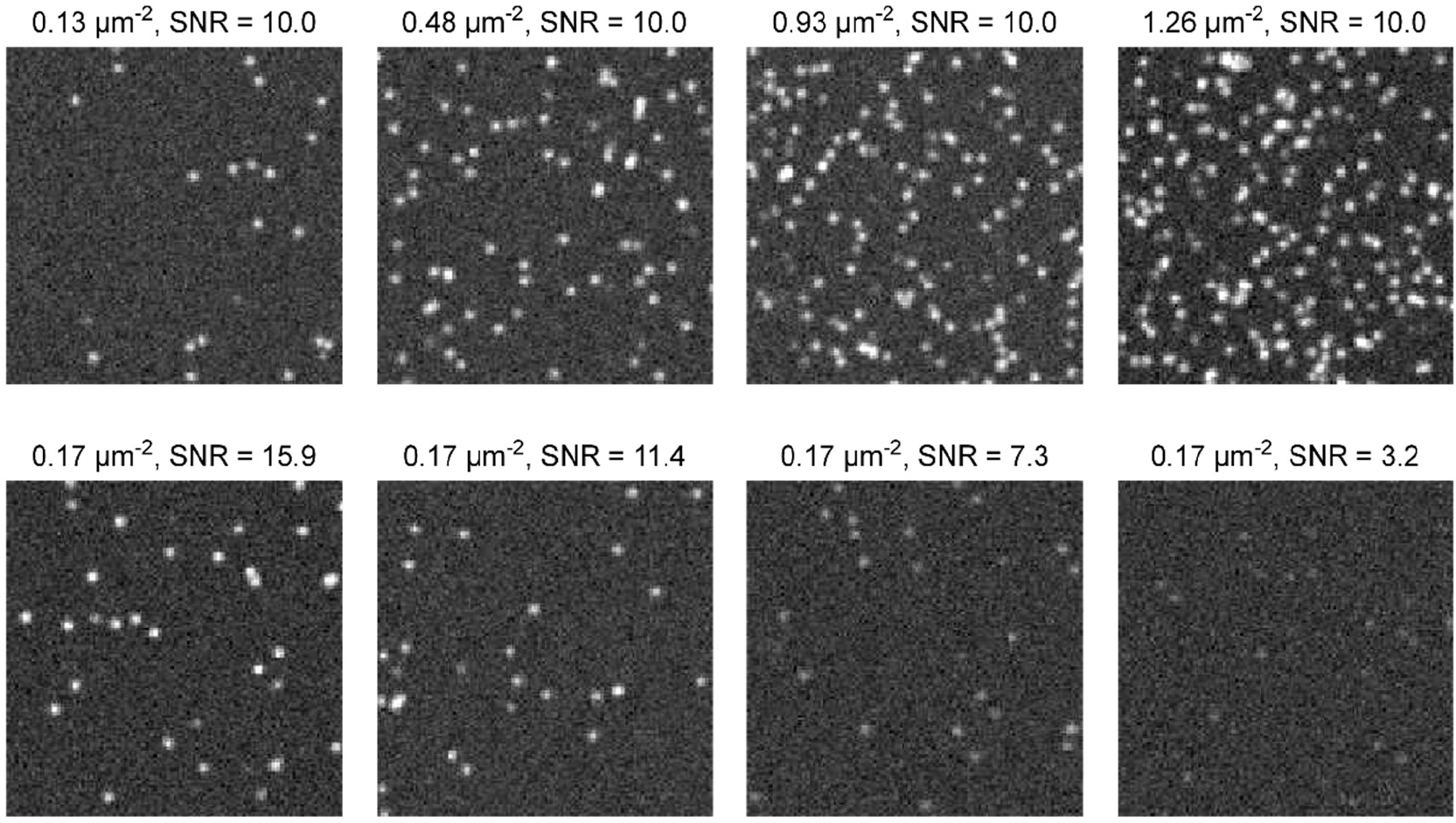
Examples of the fluorophore densities and SNRs used to test DEFCoN. The top row depicts constant SNR and increasing density from left to right; the bottom decreasing SNR and constant density.

**Supplementary Figure S4.**
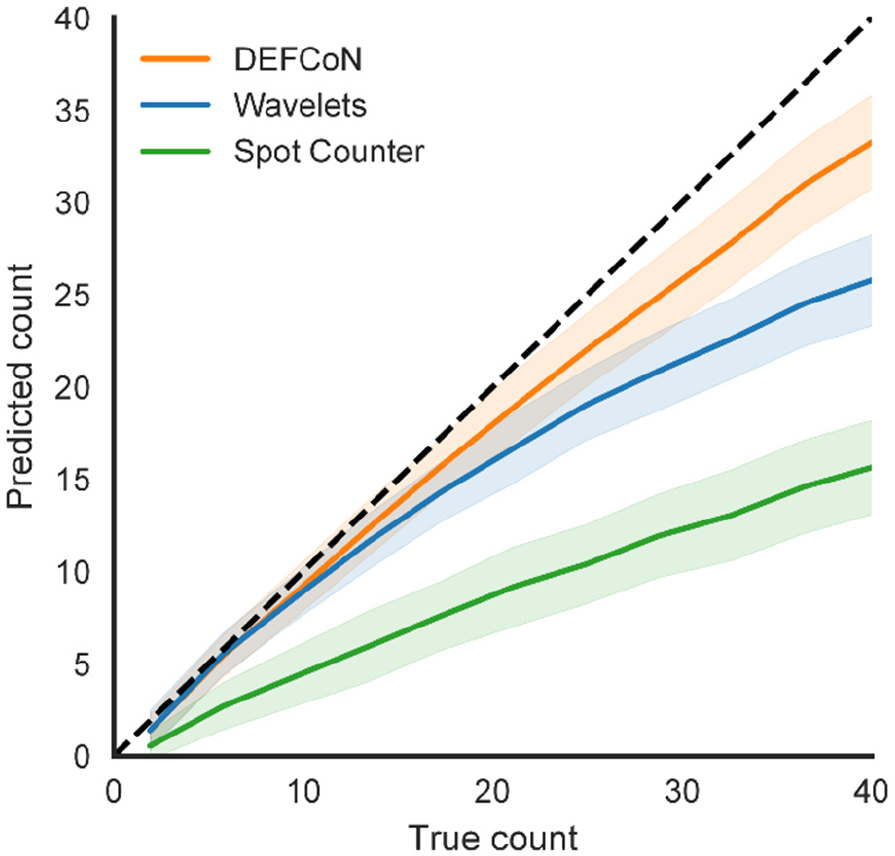
Qualitative comparison demonstrating the slower saturation of DEFCoN as compared to wavelets/watershed and a simple spot counting algorithm called Spot Counter that is based on local maxima detection. Solid lines are averages and shaded areas are standard deviation about the mean. The dashed line represents a perfect accuracy.

### E. Real dataset for comparison

Fig. S5 displays examples from the two real, i.e. not simulated, datasets from the SMLMS 2016 Challenge [1], Tubulins: Long Sequence (RealLS) and Tubulins: High Density (RealHD). Ten frames from RealLS and five from RealHD were selected and the localizations’ ground truth positions were annotated by hand. These annotations correspond to the blue circles in Fig. S5.

**Supplementary Figure S5.**
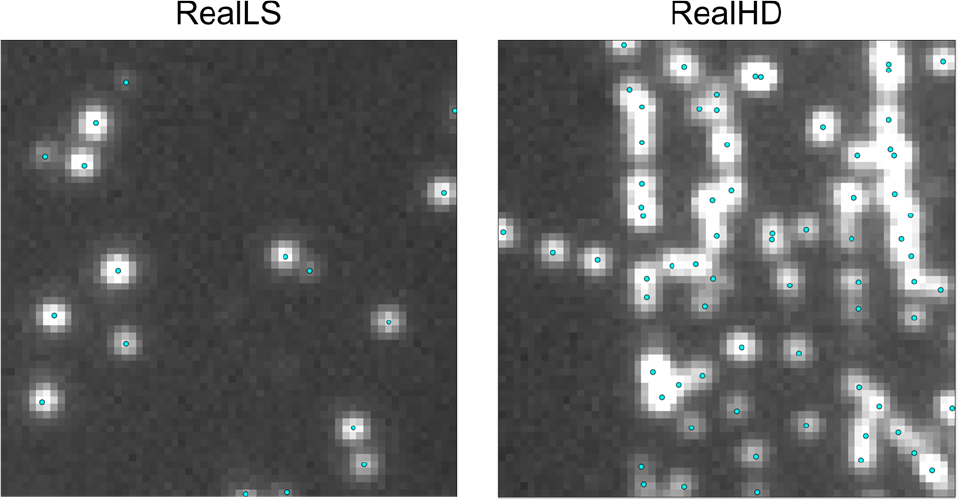
Example of annotated datasets from the SMLMS Challenge [1].

## 3. Maximum Local Count

The concept behind the maxium local count (MLC) is illustrated in Fig. S6. The sum of the pixel values over a subregion of the estimated density map returns the count estimate within that subregion. The local count around each pixel is the sum of all the pixel values from the density map within a neighborhood centered on the that pixel; convolution of the density map with a square kernel containing values all equal to one produces a local density map. The maximum value of the local density map is the MLC.

**Supplementary Figure S6.**
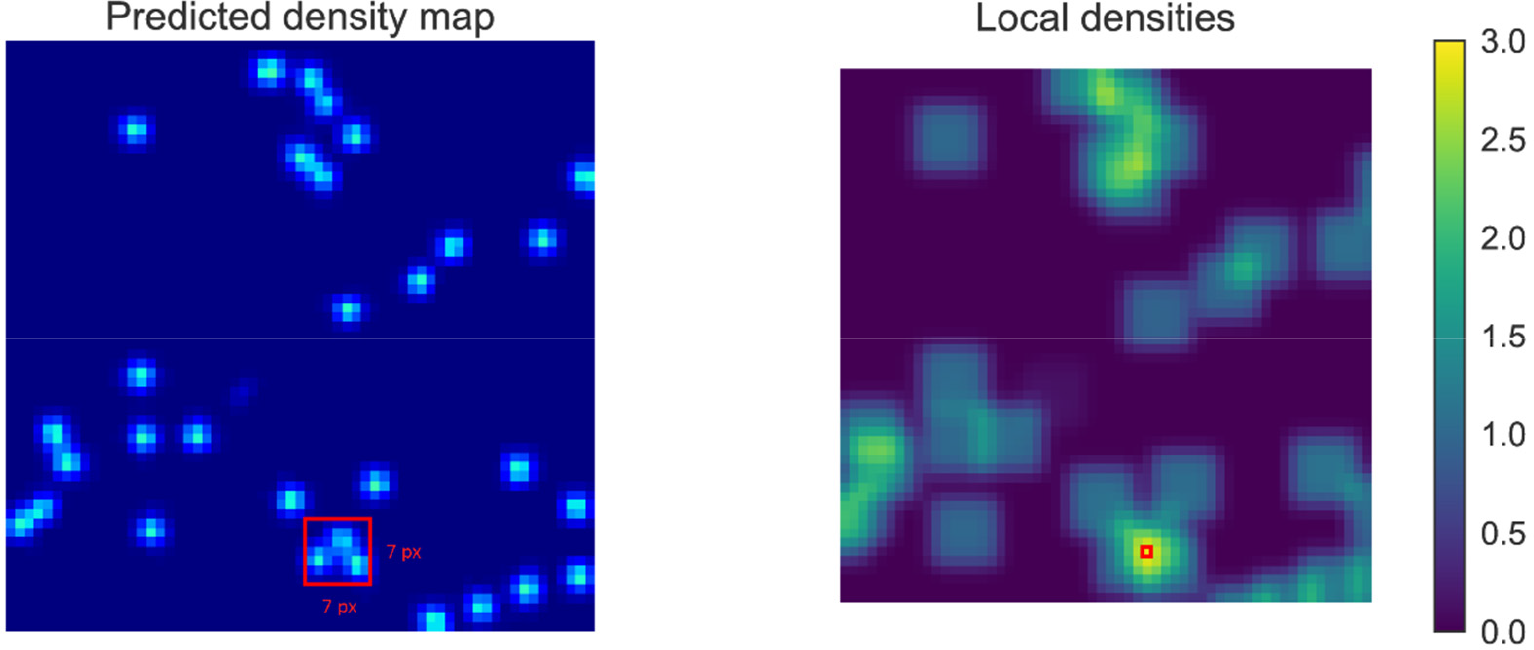
The local density estimates are made by computing the sum of the pixels in a subregion of a density map. (In this example, the size of the subregion is 7 × 7 pixels.) The maximum local count (the red square on the right) is the largest value found in the map of local density estimates.

## REFERENCES

1. M. J. Rust, M. Bates, and X. Zhuang, “Sub-diffraction-limit imaging by stochastic optical reconstruction microscopy (STORM).,” Nat. Methods 3, 793–5 (2006).

2. E. Betzig, G. H. Patterson, R. Sougrat, O. W. Lindwasser, S. Olenych, J. S. Bonifacino, M. W. Davidson, J. Lippincott-Schwartz, and H. F. Hess, “Imaging intracellular fluorescent proteins at nanometer resolution.,” Science 313, 1642–5 (2006).

3. S. T. Hess, T. P. K. Girirajan, and M. D. Mason, “Ultra-high resolution imaging by fluorescence photoactivation localization microscopy.,” Biophys. J. 91, 4258–72 (2006).

4. Y. Lin, J. J. Long, F. Huang, W. C. Duim, S. Kirschbaum, Y. Zhang, L. K. Schroeder, A. A. Rebane, M. G. M. Velasco, A. Virrueta, D. W. Moonan, J. Jiao, S. Y. Hernandez, Y. Zhang, and J. Bewersdorf, “Quantifying and optimizing single-molecule switching nanoscopy at high speeds.,” PLoS One 10, e0128135 (2015).

5. K. M. Douglass, C. Sieben, A. Archetti, A. Lambert, and S. Manley, “Super-resolution imaging of multiple cells by optimized flat-field epi-illumination,” Nat. Photonics 10, 705–708 (2016).

6. R. Diekmann, Ø. I. Helle, C. I. Øie, P. McCourt, T. R. Huser, M. Schüttpelz, and B. S. Ahluwalia, “Chip-based wide field-of-view nanoscopy,” Nat. Photonics 11, 322–328 (2017).

7. Z. Zhao, B. Xin, L. Li, and Z.-L. Huang, “High-power homogeneous illumination for super-resolution localization microscopy with large field-of-view,” Opt. Express 25, 13382 (2017).

8. A. Kechkar, D. Nair, M. Heilemann, D. Choquet, and J.-B. Sibarita, “Real-Time Analysis and Visualization for Single-Molecule Based Super-Resolution Microscopy,” PLoS One 8, e62918 (2013).

9. S. J. Holden, T. Pengo, K. L. Meibom, C. Fernandez Fernandez, J. Collier, and S. Manley, “High throughput 3D super-resolution microscopy reveals Caulobacter crescentus in vivo Z-ring organization.,” Proc. Natl. Acad. Sci. U. S. A. 111, 4566–71 (2014).

10. J. P. Eberle, W. Muranyi, H. Erfle, and M. Gunkel, “Fully Automated Targeted Confocal and Single-Molecule Localization Microscopy,” in (Humana Press, New York, NY, 2017), pp. 139–152.

11. M. Mund, J. A. van der Beek, J. Deschamps, S. Dmitrieff, J. L. Monster, A. Picco, F. Nedelec, M. Kaksonen, and J. Ries, “Systematic analysis of the molecular architecture of endocytosis reveals a nanoscale actin nucleation template that drives efficient vesicle formation,” bioRxiv 217836 (2017).

12. F. Farzam and K. A. Lidke, “Automated Multiple Target Superresolution Imaging,” in Frontiers in Optics 2017 (OSA, 2017), p. FTh3D.3.

13. A. Beghin, A. Kechkar, C. Butler, F. Levet, M. Cabillic, O. Rossier, G. Giannone, R. Galland, D. Choquet, and J.-B. Sibarita, “Localization-based super-resolution imaging meets high-content screening,” Nat. Methods 14, 1184–1190 (2017).

14. A. Burgert, S. Letschert, S. Doose, and M. Sauer, “Artifacts in single-molecule localization microscopy.,” Histochem. Cell Biol. 144, 123–31 (2015).

15. P. Fox-Roberts, R.. Marsh, K. Pfisterer, A. Jayo, M. Parsons, and S. Cox, “Local dimensionality determines imaging speed in localization microscopy,” Nat. Commun. 8, 13558 (2017).

16. J.-F. Rupprecht, A. Martinez-Marrades, Z. Zhang, R. Changede, P. Kanchanawong, and G. Tessier, “Trade-offs between structural integrity and acquisition time in stochastic super-resolution microscopy techniques,” Opt. Express 25, 23146 (2017).

17. A. D. Edelstein, M. A. Tsuchida, N. Amodaj, H. Pinkard, R. D. Vale, and N. Stuurman, “Advanced methods of microscope control using μManager software.,” J. Biol. Methods 1, e10 (2014).

18. M. Heilemann, S. van de Linde, M. Schüttpelz, R. Kasper, B. Seefeldt, A. Mukherjee, P. Tinnefeld, and M. Sauer, “Subdiffraction-resolution fluorescence imaging with conventional fluorescent probes.,” Angew. Chem. Int. Ed. Engl. 47, 6172–6 (2008).

19. D. Sage, F. R. Neumann, F. Hediger, S. M. Gasser, and M. Unser, “Automatic tracking of individual fluorescence particles: application to the study of chromosome dynamics,” IEEE Trans. Image Process. 14, 1372–1383 (2005).

20. I. Izeddin, J. Boulanger, V. Racine, C. G. Specht, A. Kechkar, D. Nair, A. Triller, D. Choquet, M. Dahan, and J. B. Sibarita, “Wavelet analysis for single molecule localization microscopy,” Opt. Express 20, 2081 (2012).

21. M. Ovesný, P. Křížek, J. Borkovec, Z. Svindrych, and G. M. Hagen, “ThunderSTORM: a comprehensive ImageJ plug-in for PALM and STORM data analysis and super-resolution imaging.,” Bioinformatics 30, 2389–90 (2014).

22. S. Holden, T. Pengo, and S. Manley, “Optimisation and control of sampling rate in localisation microscopy,” in 10th International Conference on Sampling Theory and Applications (2013), pp. 281–284.

23. “Single-Molecule Localization Microscopy: Software Benchmarking,” http://bigwww.epfl.ch/smlm/challenge2016/index.html?p=participants.

24. V. Lempitsky and A. Zisserman, “Learning To Count Objects in Images,” in Advances in Neural Information Processing Systems 23 (NIPS) (2010), pp. 1324–1332.

25. C. Arteta, V. Lempitsky, J. A. Noble, and A. Zisserman, “Interactive Object Counting,” in European Conference on Computer Vision – ECCV (Springer, 2014), pp. 504–518.

26. W. Xie, J. A. Noble, and A. Zisserman, “Microscopy cell counting and detection with fully convolutional regression networks,” Comput. Methods Biomech. Biomed. Eng. Imaging Vis. 1–10 (2016).

27. L. Fiaschi, U. Koethe, R. Nair, and F. A. Hamprecht, “Learning to count with regression forest and structured labels,” in Proceedings of the 21st International Conference on Pattern Recognition (ICPR) (2012), pp. 2685–2688.

28. D. Kang, Z. Ma, and A. B. Chan, “Beyond Counting: Comparisons of Density Maps for Crowd Analysis Tasks - Counting, Detection, and Tracking,” arXiv 1705.10118 (2017).

29. D. Oñoro-Rubio and R. J. López-Sastre, “Towards Perspective-Free Object Counting with Deep Learning,” in European Conference on Computer Vision (ECCV) (Springer, 2016), pp. 615–629.

30. L. He, X. Ren, Q. Gao, X. Zhao, B. Yao, and Y. Chao, “The connected-component labeling problem: A review of state-of-the-art algorithms,” Pattern Recognit. 70, 25–43 (2017).

31. N. Stuurman, “SpotCounter (ImageJ),” https://imagej.net/SpotCounter (2017).

32. D. Sage, H. Kirshner, T. Pengo, N. Stuurman, J. Min, S. Manley, and M. Unser, “Quantitative evaluation of software packages for single-molecule localization microscopy,” Nat. Methods 12, 717–724 (2015).

33. J. Bechhoefer, “Feedback for physicists: A tutorial essay on control,” Rev. Mod. Phys. 77, 783–836 (2005).

34. J. F. Smuts, Process Control for Practitioners: How to Tune PID Controllers and Optimize Control Loops (OptiControls Inc, 2011).

35. S. Culley, D. Albrecht, C. Jacobs, P. M. Pereira, C. Leterrier, J. Mercer, and R. Henriques, “Quantitative mapping and minimization of super-resolution optical imaging artifacts,” Nat. Methods 15, 263–266 (2018).

36. C. Allain and M. Cloitre, “Characterizing the lacunarity of random and deterministic fractal sets,” Phys. Rev. A 44, 3552–3558 (1991).

37. R. P. J. Nieuwenhuizen, K. A. Lidke, M. Bates, D. L. Puig, D. Grünwald, S. Stallinga, and B. Rieger, “Measuring image resolution in optical nanoscopy.,” Nat. Methods 10, 557–62 (2013).

38. N. Banterle, K. H. Bui, E. A. Lemke, and M. Beck, “Fourier ring correlation as a resolution criterion for super-resolution microscopy,” J. Struct. Biol. 183, 363–367 (2013).

39. T. Dertinger, R. Colyer, G. Iyer, S. Weiss, and J. Enderlein, “Fast, background-free, 3D super-resolution optical fluctuation imaging (SOFI).,” Proc. Natl. Acad. Sci. U. S. A. 106, 22287–92 (2009).

40. N. Gustafsson, S. Culley, G. Ashdown, D. M. Owen, P. M. Pereira, and R. Henriques, “Fast live-cell conventional fluorophore nanoscopy with ImageJ through super-resolution radial fluctuations,” Nat. Commun. 7, 12471 (2016).

41. E. Nehme, L. E. Weiss, T. Michaeli, and Y. Shechtman, “Deep-STORM: Super resolution single molecule microscopy by deep learning,” arxiv 1801.09631 (2018).

42. N. Boyd, E. Jonas, H. P. Babcock, and B. Recht, “DeepLoco: Fast 3D Localization Microscopy Using Neural Networks,” bioRxiv 267096 (2018).

43. M. Štefko, B. Ottino, K. M. Douglass, and S. Manley, “Design Principles for Autonomous Illumination Control in Localization Microscopy - Data,” (Version 0.1.0) [Data set]. Zenodo. http://doi.org/10.5281/zenodo.1212352 (2018).

## References

1. D. Sage, H. Kirshner, T. Pengo, N. Stuurman, J. Min, S. Manley, and M. Unser, “Quantitative evaluation of software packages for single-molecule localization microscopy,” Nat. Methods 12, 717–724 (2015).

2. S. J. Holden, T. Pengo, K. L. Meibom, C. Fernandez Fernandez, J. Collier, and S. Manley, “High throughput 3D super-resolution microscopy reveals Caulobacter crescentus in vivo Z-ring organization.,” Proc. Natl. Acad. Sci. U. S. A. 111, 4566–71 (2014).

3. N. Stuurman, “SpotCounter (ImageJ),” version 0.13, https://imagej.net/SpotCounter (2017).

4. I. Izeddin, J. Boulanger, V. Racine, C. G. Specht, A. Kechkar, D. Nair, A. Triller, D. Choquet, M. Dahan, and J. B. Sibarita, “Wavelet analysis for single molecule localization microscopy,” Opt. Express 20, 2081 (2012).

5. E. Katrukha, “ComDet,” https://github.com/ekatrukha/ComDet, (2017).

6. M. Štefko, B. Ottino, K. M. Douglass, and S. Manley, “SMLM Acquisition Simulation Software (SASS),” https://github.com/LEB-EPFL/SASS (2018).

7. S. F. Gibson and F. Lanni, “Experimental test of an analytical model of aberration in an oil-immersion objective lens used in three-dimensional light microscopy,” J. Opt. Soc. Am. A 8, 1601 (1991).

8. J. Li, F. Xue, and T. Blu, “Fast and accurate three-dimensional point spread function computation for fluorescence microscopy,” J. Opt. Soc. Am. A 34, 1029 (2017).

9. K. Perlin, “An image synthesizer,” in Proceedings of the 12th Annual Conference on Computer Graphics and Interactive Techniques - SIGGRAPH ‘85 (ACM Press, 1985), pp. 287–296.

10. K. Spencer, “Open Simplex Noise,” https://gist.github.com/KdotJPG/b1270127455a94ac5d19 (2014).

11. N. Srivastava, G. Hinton, A. Krizhevsky, I. Sutskever, and R. Salakhutdinov, Journal of Machine Learning Research: JMLR. (MIT Press, 2001), Vol. 15.

12. M. Abadi, P. Barham, J. Chen, Z. Chen, A. Davis, J. Dean, M. Devin, S. Ghemawat, G. Irving, M. Isard, M. Kudlur, J. Levenberg, R. Monga, S. Moore, D. G. Murray, B. Steiner, P. Tucker, V. Vasudevan, P. Warden, M. Wicke, Y. Yu, X. Zheng, and G. Brain, “TensorFlow: A System for Large-Scale Machine Learning,” in 12th USENIX Symposium on Operating Systems Design and Implementation (OSDI ‘16) (2016), pp. 265–284.

13. F. Chollet, “Keras,” GitHub Repos. https://github.com/fchollet/keras (2015).

14. D. P. Kingma and J. Ba, “Adam: A Method for Stochastic Optimization,” arXiv 1412.6980 (2014).

15. M. Ovesný, P. Křížek, J. Borkovec, Z. Svindrych, and G. M. Hagen, “ThunderSTORM: a comprehensive ImageJ plug-in for PALM and STORM data analysis and super-resolution imaging.,” Bioinformatics 30, 2389–90 (2014).

